# Broad H3K4me3 Domain Is Associated with Spatial Coherence during Mammalian Embryonic Development

**DOI:** 10.1101/2023.12.11.570452

**Authors:** Xuan Cao, Terry Ma, Rong Fan, Guo-Cheng Yuan

## Abstract

It is well known that the chromatin states play a major role in cell-fate decision and cell-identity maintenance; however, the spatial variation of chromatin states *in situ* remains poorly characterized. Here, by leveraging recently available spatial-CUT&Tag data, we systematically characterized the global spatial organization of the H3K4me3 profiles in a mouse embryo. Our analysis identified a subset of genes with spatially coherent H3K4me3 patterns, which together delineate the tissue boundaries. The spatially coherent genes are strongly enriched with tissue-specific transcriptional regulators. Remarkably, their corresponding genomic loci are marked by broad H3K4me3 domains, which is distinct from the typical H3K4me3 signature. Spatial transition across tissue boundaries is associated with continuous shortening of the broad H3K4me3 domains as well as expansion of H3K27me3 domains. Our analysis reveals a strong connection between the genomic and spatial variation of chromatin states, which may play an important role in embryonic development.

## Introduction

The dynamic change of chromatin states provides an important layer of gene regulatory mechanism that enables cells to respond to developmental cues, environmental signals, and various physiological demands ^1,2^. The development of genome-wide assays at increasingly refined cell resolution has greatly enabled biologists to characterize chromatin dynamics and to investigate its biological function ^3–12^. On the other hand, few studies have examined the spatial context of chromatin dynamics ^13–18^. As such, the role of the tissue environment in mediating chromatin dynamics remains poorly understood.

Recently, several technologies have been developed to profile genome-wide chromatin states while maintaining the spatial information ^11,13–18^, thereby providing a great opportunity to systematically characterize the tissue-wide spatial organization of chromatin states and to further investigate the underlying mechanisms. On the other hand, previous analyses of such data have been limited to spatial mapping of various cell-types, whereas the relationship between spatially adjacent cell neighborhoods has not been systematically investigated. As such, a coherent view of the tissue-level structural organization remains lacking.

To fill this gap, we have adapted recently developed spatial transcriptomics ^19,20^ and chromatin state analysis tools ^21,22^ to systematically characterize the tissue-level spatial organization of chromatin state profiles in the mouse embryo ^13^. Through these analyses, we identified spatially coherent chromatin state patterns that recapitulated the anatomic structure, and found that the associated genes comprised many tissue-specific transcriptional regulators that are important for organ development. Strikingly, the genomic loci of the spatially coherent genes are distinctly marked by broad domains of H3K4me3, which typically has a narrow peak signature. Our analysis reveals a previously unrecognized link between the genomic and spatial organization of chromatin states, and suggests that such a link may play an important role in embryonic development.

## Results

### Spatial-CUT&Tag data analysis identified distinct cell clusters in the mouse embryo

To characterize the spatial organization of histone modification and investigate its role in embryonic development, we systematically analyzed a recently published spatial-CUT&Tag dataset derived from embryonic day 11 (E11) mouse embryo ^13^. Dimensionality reduction and clustering of H3K4me3 by using ArchR ^21^ partitioned the 2122 spots into 9 distinct clusters spanning the E11 mouse embryo (Figures S1A and S1B). We manually annotated clusters based on the activities of known marker genes (Figures S1C and S1D; STAR Methods). For example, the heart cluster corresponds to high activities of Gata4 and Gata6 ^21,23,24^, which are essential for regulating the onset of cardiac myocyte differentiation during mammalian development. The spots from the forebrain cluster are marked with strong H3K4me3 of gene Foxg1 ^25^, which plays critical roles in forebrain development. The meninges cluster is marked by the high activity of Foxc2 ^26^, which is a key factor in meningeal lymphatic vessels. These annotations are further validated by visual inspection of their respective anatomic locations (Figure S1B). Several clusters are not clearly associated with specific marker genes. To annotate these clusters, we only used the spatial information.

### Spatially coherent H3K4me3 patterns are strongly linked to the structural organization of the mouse embryo

Since clustering methods do not consider spatial information, we applied the BinSpect (Binary Spatial extract) method from the Giotto package to identify spatial genes ^19^. Among the top 200 identified spatial genes, several are well-known tissue-specific regulators, such as Gata6 in heart and Pou3f2 (also known as Oct7) in three brain regions (forebrain, midbrain, and hindbrain) (Figure S1D; Table S1) ^19,24,27^, suggesting that the spatially coherent distribution of H3K4me3 at key genes are related with cell lineage maintenance.

To systematically investigate the spatial patterns and biological functions of the identified spatial genes, we divided them into 8 distinct modules based on spatial co-localization analysis (Figure 1A; STAR Methods; Table S1)^19^. For each module, we summarized the overall spatial pattern by averaging the H3K4me3 scores associated with all its component genes (Figure 1A). These patterns correspond well to the underlying anatomic structures, and display clearer spatial continuity than the cell clusters identified in the previous section (Figure 1A). To systematically characterize the biological functions of the spatial genes in each module, we performed functional enrichment analysis based on gene ontology (GO) annotations ^28^. The most enriched terms are all related to organism development (Figure 1B). To precisely map the modules to cell clusters, we performed gene set enrichment analysis using the PAGE (Parametric Analysis of Gene Set Enrichment) algorithm ^29^. We found that a co-localization module is mapped either to a unique cell cluster or to multiple functionally related and spatially adjacent clusters (Figures 1C and 1D). For example, Module M1 primarily maps to the heart, although it is also slightly enriched in the surrounding cartilage ^30^. Module M2 contains the largest number of spatial genes (n = 84) (Table S1), and it is uniquely mapped to the liver. Module M3 is enriched in three brain regions (forebrain, midbrain, and hindbrain) with no obvious biases. In contrast, Module M4 is specifically enriched at the forebrain region. Module M5 maps to multiple anatomic regions including the facial and limb, meninges and the spinal cord. M6 is uniquely mapped to the limb. Module M7 maps to cartilage, a partial area of spinal cord and hindbrain. Lastly, Module M8 maps to the spinal cord.

**Figure 1.**
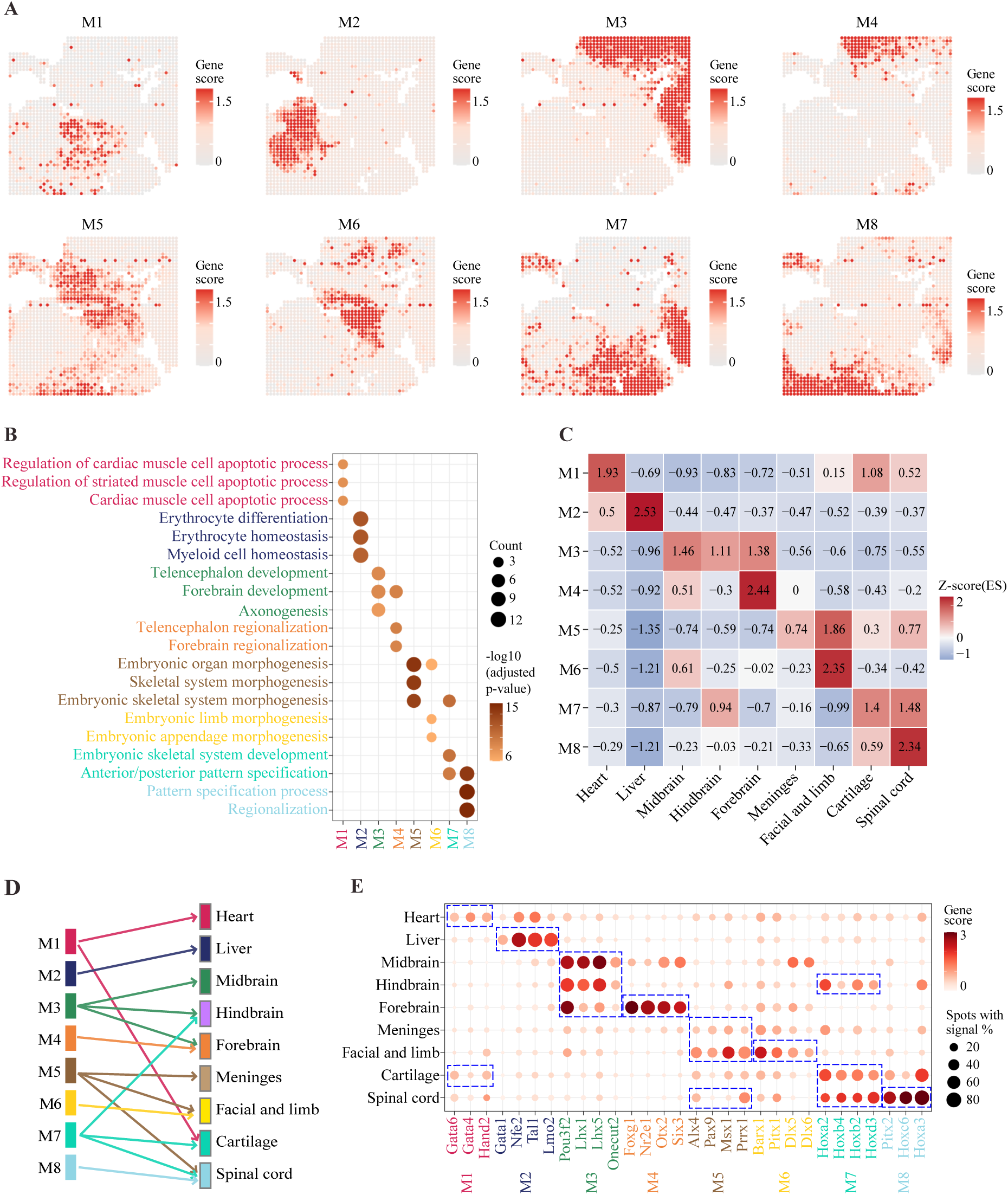
Identification of spatial genes and colocalization modules. (A) The average spatial pattern for identified 8 distinct H3K4me3 colocalization modules (M1-M8). (B) The three most significantly enriched GO terms associated with each colocalization module. (C-D) Mapping colocalization modules to annotated clusters. (C) Using gene set enrichment analysis to quantify the degree of association between colocalization modules and clusters. The raw enrichment scores are converted to z-scores for comparison. (D) The discretized mapping between modules and clusters resulting from using z-score = 0.7 as the cutoff. (E) Clusters specific H3K4me3 signals of the top spatially coherent genes in each colocalization module.

Of note, tissue-specific transcription factors (TFs) are over-represented in the colocalization modules (Figure 1E). Module M1, which corresponds to the heart cluster, contains TFs Gata6, Gata4 and Hand2, which are essential TFs in cardiomyocyte differentiation ^23,24,31^. The liver associated module M2 contains master regulators for hematopoiesis, such as Tal1 and Lmo2 ^32–34^. The brain associated module M3 contains several TFs that are known to play essential roles in brain development, such as Pou3f2, Lhx1, and Lhx5 ^27,35^. Several regulators contained in the module M4 are highly associated with the development of the forebrain, such as Foxg1 and Nr2e1 ^36,37^. The module M5 contains several TFs that are related to limb development (such as Alx4 and Prrx1 ^38,39^) and craniofacial and tooth development (such as Msx1 and Pax9 ^40,41^). M6, whose spatial pattern is similar to M5, also contains TFs that are related to limb and craniofacial, such as Dlx5, Dlx6, Barx1, and Pitx1^42–44^. M7 contains multiple Hox family TFs that are involved in body patterning, including Hoxa2, Hoxb2, and Hoxb4 ^45^. Taken together, these analyses strongly suggest that the maintenance of spatial coherence of H3K4me3 profiles may play an important role in shaping the tissue structural organization during embryonic development.

#### Spatial genes are characterized by broad domains

We examined the genomic landscape of the H3K4me3 signal to search for distinct properties associated with the spatial coherence. To this end, we constructed pseudo bulk data for each cell cluster by averaging the H3K4me3 signal over all spots that belong the cluster. To systematically annotate chromatin states at multiple scales, we applied the diHMM model ^22^, which simultaneously provides both nucleosome- and domain-level state annotations. This approach allows us to clearly distinguish narrow peaks from broad domains in a common model framework. Following previous studies ^46,47^, we used a 5kb width cutoff to identify broad domains. Strikingly, we found most spatial genes, including all the examples mentioned above, are marked by broad H3K4me3 domains at their promoter regions in the corresponding cellular context (Table S2). For example, the Gata6 locus is marked by a 11.5kb wide broad domain in the heart cluster (Figure S2A; Table S2). Similarly, the Pou3f2 locus is marked by a 22 kb wide broad domain in the brain, and the Foxg1 locus is marked by a 21.5kb wide broad domain in the forebrain (Figure 2A; Figure S2A; Table S2). In contrast, housekeeping genes, such as Copb2 ^48^ are typically associated with narrow peaks (domain width 1-2kb) (Figure 2A), although a few counter-examples exist (such as Actb). Of note, the broad domains are formed specifically in those genes’ spatially defined cellular context (Figure 2B).

**Figure 2.**
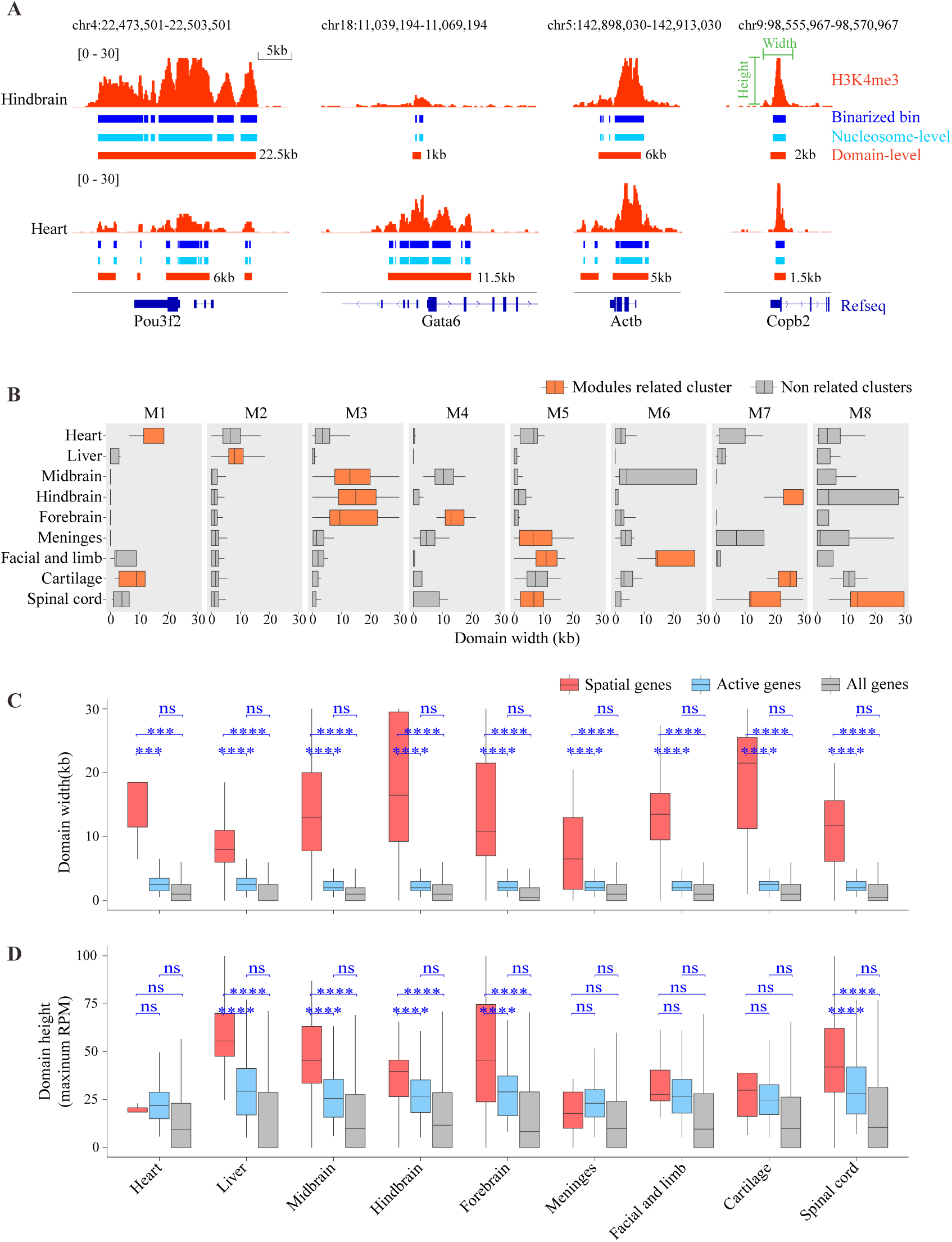
Broad H3K4me3 domains mark spatial genes. (A) Genome browser tracks showing H3K4me3 raw and binarized signals along with the diHMM output: nucleosome- and domain-level chromatin state annotation at representative spatial genes (Pou2f2 and Gata6) and housekeeping genes (Actb and Copb2). (B) The distribution of domain width at the genomic loci of the spatially coherent genes. (C-D) Comparisons of the H3K4me3 domain width (C) and height (D) between spatial genes, activate genes, and all genes (genome-wide) in each cluster. Spatial genes in each cluster are spatial genes identified from related modules as in 2B and 2E. Activate genes are genes that promoter regions occupancy by H3K4me3 domains excluding the spatial gene. ns (not significant) indicate p-value > 0.05 using Wilcoxon rank-sum test; *** indicate p-value ≤ 0.001; **** indicate p-value ≤ 0.0001.

To evaluate the overall degree of association between spatial genes and broad domains, we compared the domain width of spatially coherent genes (n = 5 - 84), active genes (n = 11,000 - 13,000), and genome-wide genes (n = 23,995) to measure the H3K4me3 domain width (Figures S2B and S2C). Spatially genes are associated with broader domains (median domain width 6.5kb - 21.5kb) compared to other active genes (median domain width 2kb - 2.5kb) and genomic background (median domain width 0kb - 1kb) (Figure 2C; Figure S2D). In comparison, the difference in domain height, quantified as the maximal H3K4me3 signal level within each domain, is modest (Figure 2D; Figure S2D). As such, we have discovered a strong association between broad domains and spatial genes.

#### Distinct spatial profile of H3K27me3 in E11 mouse embryo

Tissue specific changes of H3K4me3 are mediated by its dynamic interplay with other histone marks. To explore the spatial relationship between H3K4me3 and other histone marks, we focused on H3K27me3, which is a repressive histone mark ^47,49,50^. In their original study ^13^, the authors also generated the spatial H3K27me3 profile in an adjacent tissue slice obtained from the same mouse embryo. To integrate H3K4me3 and H3K27me3 profiles, we aligned the images by identifying the orthogonal transformation that maximizes the intensity gradient across embryo boundaries using an iterative algorithm (Figure S3A; STAR Methods). After alignment, we transferred the measured H3K27me3 profiles to the overlapping region, which contains 1571 of the 2122 spots whose H3K4me3 profiles had been analyzed (Figure 3A; Figure S3B). Due to the lack of overlap, we removed the spinal cord cluster from further analysis.

**Figure 3.**
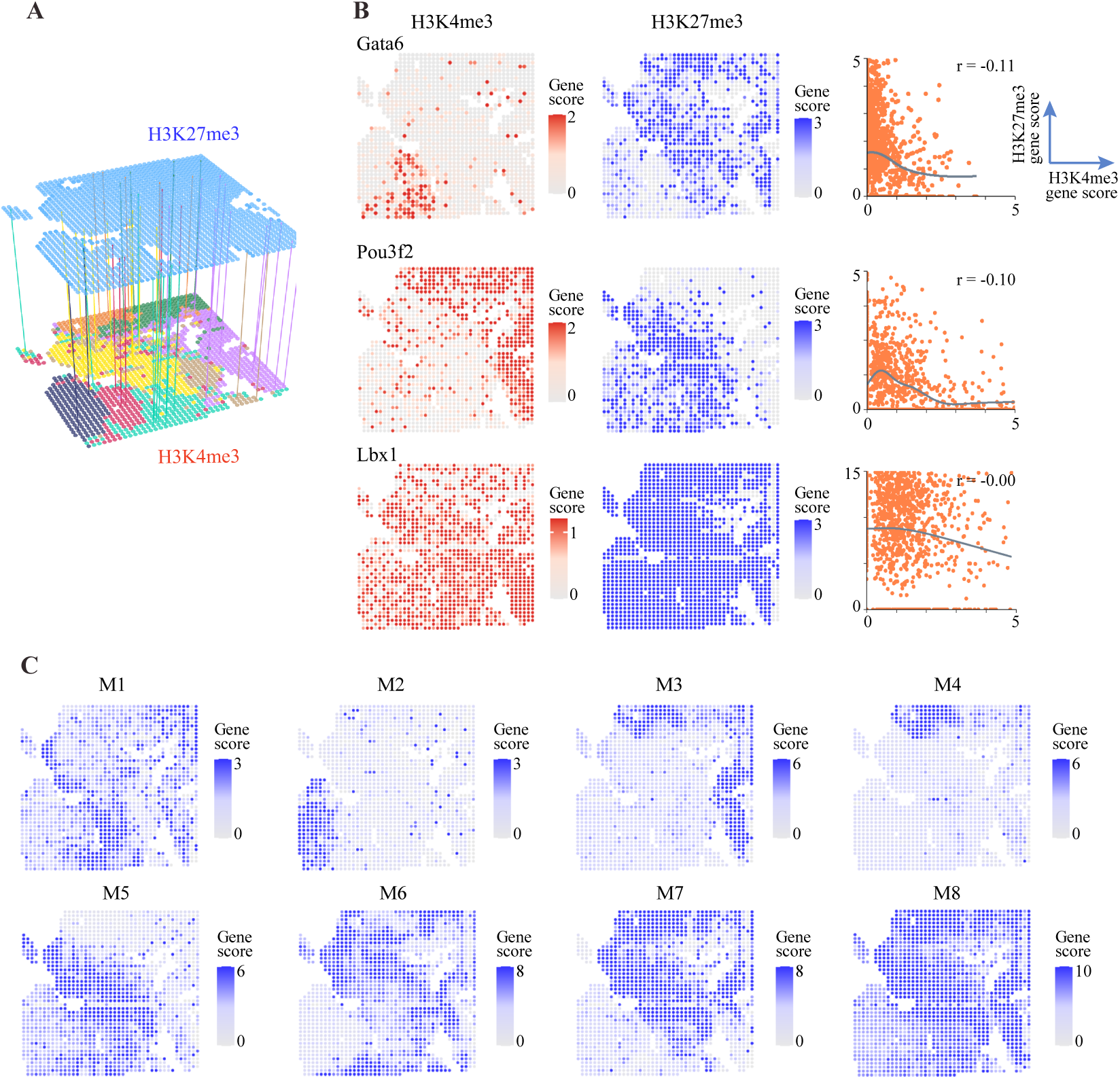
Integrated analysis of H3K4me3 and H3K27me3 data. (A) A schematic view of our approach to align H3Kem3 and H3K27me3 data. (B) The relationship between H3K4me3 and H3K27me3 patterns. The spatial distribution of H3K4me3 (left) and H3K27me3 (middle) signals. Scatter plot (right) shows different patterns of correlation between H3K4me3 and H3K27me3 signals. r is the pearson correlation; black lines are fitted curves by loess regression. (C) The average spatial pattern for identified 8 distinct H3K27me3 colocalization modules (M1-M8).

We observed several types of relationship between the spatial patterns of H3K27me3 and H3K4me3 associated with a gene (Figure 3B; Figure S3C). For example, in the heart, where Gata6 has strong signals, the H3K27me3 signal is nearly absent. Outside the heart, the H3K4me3 signal is greatly diminished, replaced by elevated H3K27me3. Similarly, the H3K4me3 signal for Pou3f2 is concentrated in the brain, whereas H3K27me3 shows elevated activity in other regions patterns (Figure 3B). Such a complementary relationship between H3K4me3 and H3K27me3 is most common. At the extreme, for housekeeping genes, such as the Actb ^48^, the entire embryo is marked with H3K4me3 activity at the absence of H3K27me3, whereas Hoxc9 is marked only by H3K27me3 across the entire embryo (Figure S3C). However, there exist other types of spatial relationship. For example, Gata1 has a strong H3K4me3 signal in the liver, but it is not replaced by H3K27me3 outside of the liver (Figure S3C). In addition, the co-occupancy of H3K4me3 and H3K27me3, the “bivalent mark”, was enriched at several genes, such as Lbx1 ^51^ (Figure 3B).

By applying the spatial gene analysis to H3K27me3, we identified 200 spatial genes and 8 co-localization modules (Figure 3C; Figure S3D; Table S3). Unlike H3K4me3, a H3K27me3 module typically spans across multiple tissues (Figure 3C). For example, both M7 and M8 encompass most cell clusters. These modules contain multiple Hox family genes whose activities may guide the spatial patterning during embryonic development ^52,53^. Among the top 200 spatial genes, 39 are shared between H3K4me3 and H3K27me3 analysis (Table S4). For these genes, the spatial patterns of H3K4me3 and H3K27me3 are typically complementary.

#### Spatial transition of chromatin states across tissues

To further investigate the connection between genomic and spatial variation of chromatin states, we were interested to examine how the genomic tracks of H3K4me3 vary spatially across tissue boundaries. To reduce the impact of the variation of tissue morphologies, we defined a spatial pseudo distance metric that takes into account both H3K4me3 signal and spatial differences (Figure 4A; STAR Methods). Using this metric, we were able to quantify the pseudo distance between each spot to the center of any tissue of interest, such as the heart (Figure 4B). To enhance robustness, we further divided spots into 20 equidistance groups with increasing order. For each group, we averaged the H3K4me3 profile and examined the domain width corresponding to the spatial genes.

**Figure 4.**
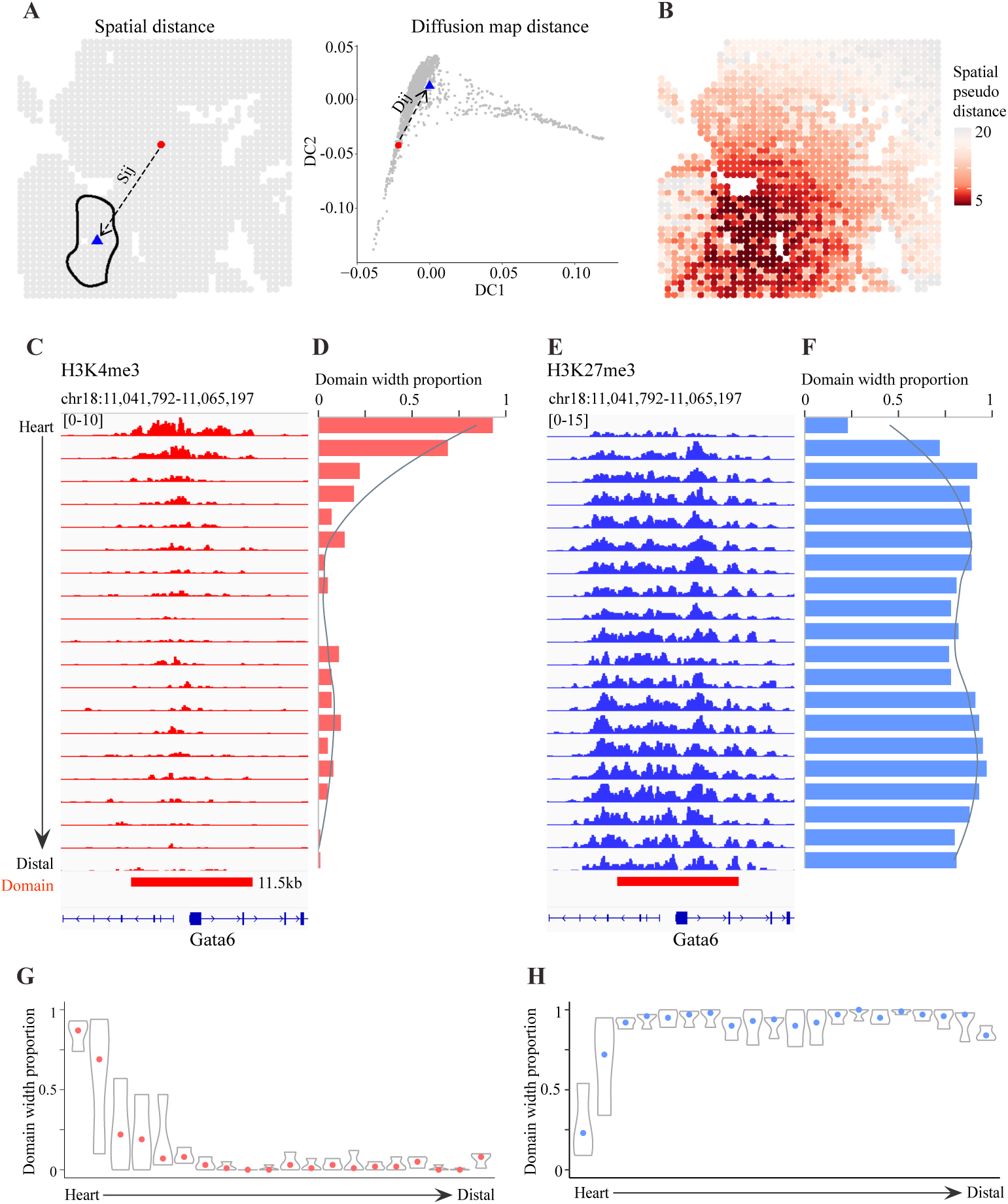
Spatial transition of chromatin states across tissues (using heart as a representative example) (A) A schematic illustration of the spatial pseudo distance between a source (red circle) and target (blue triangle) spot. It incorporates both physical distance (S_ij_) and chromatin state similarity (D_ij_). (B) The spatial distribution of the spatial pseudo distance to the center of the heart. (C) Genome browser tracks showing H3K4me3 signal at Gata6 in each equidistance group from the heart center. (D) The H3K4me3 domain width corresponds to each equidistance group. (E) Similar to (C) but for H3K27me3. (F) Similar to (D) but for H3K27me3. (G) The distribution of domain width for all spatial genes in the M1 H3K4me3 module. (H) Similar to (G) but for H3K27me3.

We first examined the pattern of spatial transition from the heart to surrounding tissues, using Gata6 as a representative example. As noted above, the H3K4me3 signal at Gata6 is highly specific to the heart, and its corresponding locus displays a broad domain pattern. By examining the change of domain width with respect to the spatial pseudo distance to the center of the heart, we found that the domain width at Gata6 decreases continuously across the heart boundary (Figures 4C and 4D). Due to the sparsity of the data, it is unclear whether this reflects coordinated change of chromatin states or increased cellular heterogeneity. Nonetheless, this observation is consistent with the view that the formation of a broad domain is a complex process involving many factors ^54^. To gain insights into a potential role of H3K27me3 in mediating the width of H3K4me3 domains, we repeated the above analysis for the H3K27me3 signal. Here we observed an opposite trend, with the H3K27me3 domain width increasing across the heart boundary (Figures 4E and 4F).

We repeated this analysis to other genes and tissues and found a similar pattern (Figures 4G and 4H; Figure S4). In general, tissue-specific broad domains shorten continuously from the center of the tissue. While in most cases, the shortening of a H3K4me3 domain is accompanied by the broadening of the corresponding H3K27me3 domain, there are also notable exceptions such as the liver (Figure S4). Taken together, these data suggest spatially continuous shortening of broad H3K4me3 domains may play an important role for organism patterning. This process is often mediated in coordination with H3K27me3 pattern changes.

## Discussion

Spatial biology has provided powerful tools to map the inner structure of tissues and organs and to investigate the role of tissue microenvironment in mediating cell states. Previous analyses based on gene expression profiles have identified distinct cell neighborhoods and spatial domains, whereas the emergence of spatial epigenomic technologies offers an opportunity to investigate the underlying mechanisms. By using recently developed spatial omic and chromatin state analysis tools, we were able to gain new insights that may guide future mechanistic investigations. In particular, we showed that many tissue-specific regulators have spatially coherent chromatin state patterns. Of note, the target genes of these regulators typically do not share similar spatial patterns, suggesting that the maintenance of spatial coherence is a distinct property of tissue-specific regulators. Furthermore, we found that the spatially coherent genes are distinctly marked by H3K4me3 broad domains, suggesting that the establishment of broad domains may serve as an important mechanism for maintaining spatial coherence. Our analysis is consistent with and further extends previous findings that broad H3K4me3 domains play an important role in cell identity maintenance ^55,56^. Further studies are needed to study the mechanisms underlying their formation and disassociation, as well as the role of tissue microenvironment in mediating such changes. H3K4me3 broad domains share similar properties with super-enhancers which have been intensively studied in recent years ^57^. It would be interesting to study whether and how two marks spatially change in coordination. Unfortunately, we were not able to explore this direction due to the lack of matching H3K27ac data.

It is well-known that antagonistic activities of H3K4me3 and H3K27me3 play an important role in gene regulation, and that bivalent domains are signatures of developmental regulators^51^. The spatial distributions of H3K4me3 and H3K27me3 are often complementary (Figure 3B; Figure S3C), suggesting that the competition between H3K4me3 and H3K27me3 activities may play a role in shaping the spatial transition of chromatin states. It will be interesting to investigate whether there are distinct chromatin regulators that mediate such transitions.

We recognize that spatial epigenomics analysis is only emerging. New computational tools are needed to effectively utilize the spatial information in such data to build gene regulatory networks that can incorporate both cell-type intrinsic regulators and cell-cell interaction mechanisms. Furthermore, emerging technologies can profile multiple data modalities from the same cells while maintaining spatial information. Such data will be extremely useful both for comprehensive cell-state mapping and for investigating the underlying mechanisms. Moving forward, we envision that the development of new computational and experimental methods of spatial-omics datasets will continue to allow us to explore regulatory relationships and how distinct cell states respond to development and diseases.

## Acknowledgments

We would like to thank Yanxiang Deng at University of Philadelphia and Di Zhang at Yale University for sharing their datasets and insightful discussions. This research was supported by National Institutes of Health grants (RF1MH133703 to G.-C.Y. and RF1MH128970 to G.-C.Y.).

Author Contributions

G.-C.Y. conceived and supervised the project. X.C. and G.-C.Y. conducted and supervised the computational analyses. X.C. T.M. helped with image alignment. X.C. and G.-C.Y. wrote the manuscript with input from R.F. All of the authors contributed ideas for this work. All of the authors reviewed and approved the manuscript.

## STAR Methods

### Key Resources Table

| REAGENT or RESOURCE | SOURCE | IDENTIFIER |
| --- | --- | --- |
| **Deposited data** |  |  |
| Raw spatial-CUT&Tag of H3K4me3 and H3K27me3 datasets | Deng et al.<sup>13</sup> | https://www.ncbi.nlm.nih.gov/geo/query/acc.cgi?acc=GSE165217 |

| Software and algorithms |  |  |
| --- | --- | --- |
| R (v4.1.3) | The R Foundation | https://www.r-project.org |
| ArchR (v1.0.1) | Granja et al.<sup>21</sup> | https://www.archrproject.com |
| Giotto (v1.1.2) | Dries et al.<sup>19</sup> | https://github.com/drieslab/Giotto |
| diHMM | Kai et al.<sup>58</sup> | https://github.com/ykai16/diHMM |
| bedtools (v2.29.2) | Quinlan et al.<sup>59</sup> | https://github.com/arq5x/bedtools2 |
| IGV (v2.12.3). | Robinson et al.<sup>60</sup> | https://software.broadinstitute.org/software/igv/ |
| clusterProfiler (v4.2.2) | Yu et al.<sup>61</sup> | http://bioconductor.org/packages/release/bioc/html/clusterProfiler.html |
| destiny (v3.8.1) | Angerer et al.<sup>61,62</sup> | https://github.com/theislab/destiny |

### Resource availability

#### Lead Contact

Further information and requests for resources should be directed to and will be fulfilled by the Lead Contact, Guo-Cheng Yuan.

### Method details

#### Pre-processing spatial-CUT&Tag data

We used ArchR (v1.0.1) ^21^ to pre-process the data and perform downstream analyses of the spatial-CUT&Tag data ^13,21^. We converted the fragments file into a tile matrix in 5kb genome binning size (based on mm10) and pixels not on tissue were removed based on the metadata file. The data were normalized using iterative latent semantic indexing (LSI) (iterations = 3, resolution = 0.2, varFeatures = 25000, dimsToUse = 1:30, sampleCells = 10000, n.start = 10) and clustered with a resolution of 1. Uniform Manifold Approximation and Projection (UMAP) ^63^ embeddings performed based on LSI (nNeighbors = 20, minDist = 0.4, metric = cosine). Gene score matrices were calculated using the Gene Score model in ArchR.

#### Spatially coherence genes detection

We applied Giotto (v1.1.2) ^19^ to evaluate spatially coherent genes. Firstly, we loaded the gene score matrix, together with the spatial coordinates of the spots, to create the *giotto* object. Secondly, we created the spatial network connecting spots based on their physical distance by applying the *createSpatialNetwork* function (method = kNN, minimum_k = 0, k=10, maximum_distance_knn = 40). To calculate spatially coherent genes, we used the *binSpect* (Binary Spatial extract) function and selected top 200 genes ranked by adj.p.value in further analysis.

#### Spatial co-localization modules

To identify spatial co-localization modules, we applied the binSpect method implemented in Giotto and identified the top 200 spatially coherent genes. We used the function *detectSpatialCorGenes* (method = network, spatial_network_name = kNN_network) to calculate a gene-to-gene correlation score matrix followed by function *clusterSpatialCorGenes* (hclust_method = ward.D2, k = 8) to identified co-localization modules by hierarchical clustering the gene-to-gene correlation score matrix. Finally, we used the function *createMetagenes t*o summarize the overall spatial pattern for each module.

#### Mapping co-localization modules to cell clusters

To map each colocalization module to the cell types identified by clustering analysis, we compute an enrichment score by using the PAGE (Parametric Analysis of Gene Set Enrichment) ^29^ analysis. Briefly, we calculate the foldchange of gene activity scores compared for each spatial gene within a module. Then we computed the enrichment scores based on the mean and standard deviation of the fold change values. The enrichment score values are then normalized by converting to z-scores.

#### Multi-scale chromatin state annotation by using diHMM

We applied the diHMM method to identify both nucleosome- and domain-level chromatin states, using its Python/C++ implementation ^58^. The fragments of H3K4me3 across each cluster were divided into 100 bp bins, and the fragment counts were binarized by using n=5 cutoff. Next, we trained the diHMM model by using the following parameter setting: domain_size=5, domain_states=2, bin_states=2, bin_size=100.

#### Function enrichment analysis

Function enrichment analyses were performed using clusterProfiler (v4.2.2) ^61^. The identified *biological_process* Gene Ontology (GO) terms were ranked by their associated adjusted p-values. The top three terms for each cluster were reported.

#### Integration of H3K4me3 and H3K27me3 data

The Iterative Closest Points (ICP) algorithm ^64^ is used to align the tissue-slice microscopy images corresponding to the H3K4me3 and H3K27me3 assays. Briefly, the imaging data are first converted to point clouds by using the DBSCAN algorithm ^65^. Each point corresponds to the centroid of a cluster identified by DBSCAN. For each point in the source point cloud (corresponding to H3K27me3), we find the closest point in the target point cloud (corresponding to H3K4me3) using a nearest-neighbor search. This creates pairs of corresponding points. We then used singular vector decomposition to identify the optimal transformation that preserves the covariance matrix associated with the point cloud mapping. These two steps are iteratively refined until convergence. The aforementioned procedure was used to align the whole embryo images, and then restricted to spatial-CUT&Tag field of views (FOVs). Based on the alignment result, we inferred the H3K27me3 gene score values at matched spots by using numerical interpolation.

#### Spatial pseudo distance calculating

To assist the investigation of spatial transition, we defined a spatial pseudo distance to account for both chromatin state and physical distance differences, as follows. For each pair of points i and j, we use S_ij_ to represent their physical distance (Figure 5A, left). To quantify their chromatin state difference, we first projected the high-dimensional H3K4me3 gene score data to a 2D space (DC1 and DC2) by using the diffusion maps, as implemented in the *destiny* package ^62,66^. Then we used the Euclidean distance D_ij_ in this reduced space to quantify the chromatin state differences (Figure 5A, right). The spatial pseudo distance is then defined as the sum of the normalized S_ij_ and D_ij_ values. To measure the deviation from each tissue (e.g., heart), we evaluated the spatial pseudo distance between each spot and the centroid of the tissue. The spots are divided into 20 equidistance groups to enhance the robustness of the analysis. For each broad domain in question, we determined its width in each group of spots by counting the fraction of areas that have significant H3K4me3 or H3K27me3 signals.

## Supplemental information

Supplemental figures S1-S4.

Table S1. Top 200 H3K4me3 spatial genes for each colocalization module (M1-M8), related to Figure 1.

Table S2. Spatial genes with broad H3K4me3 domain, related to Figure 2.

Table S3. Top 200 H3K27me3 spatial genes for each colocalization module (M1-M8), related to Figure 3.

Table S4. Shared spatial genes between H3K4me3 and H3K27me3, related to Figure 3.

---

**Figure S1.**
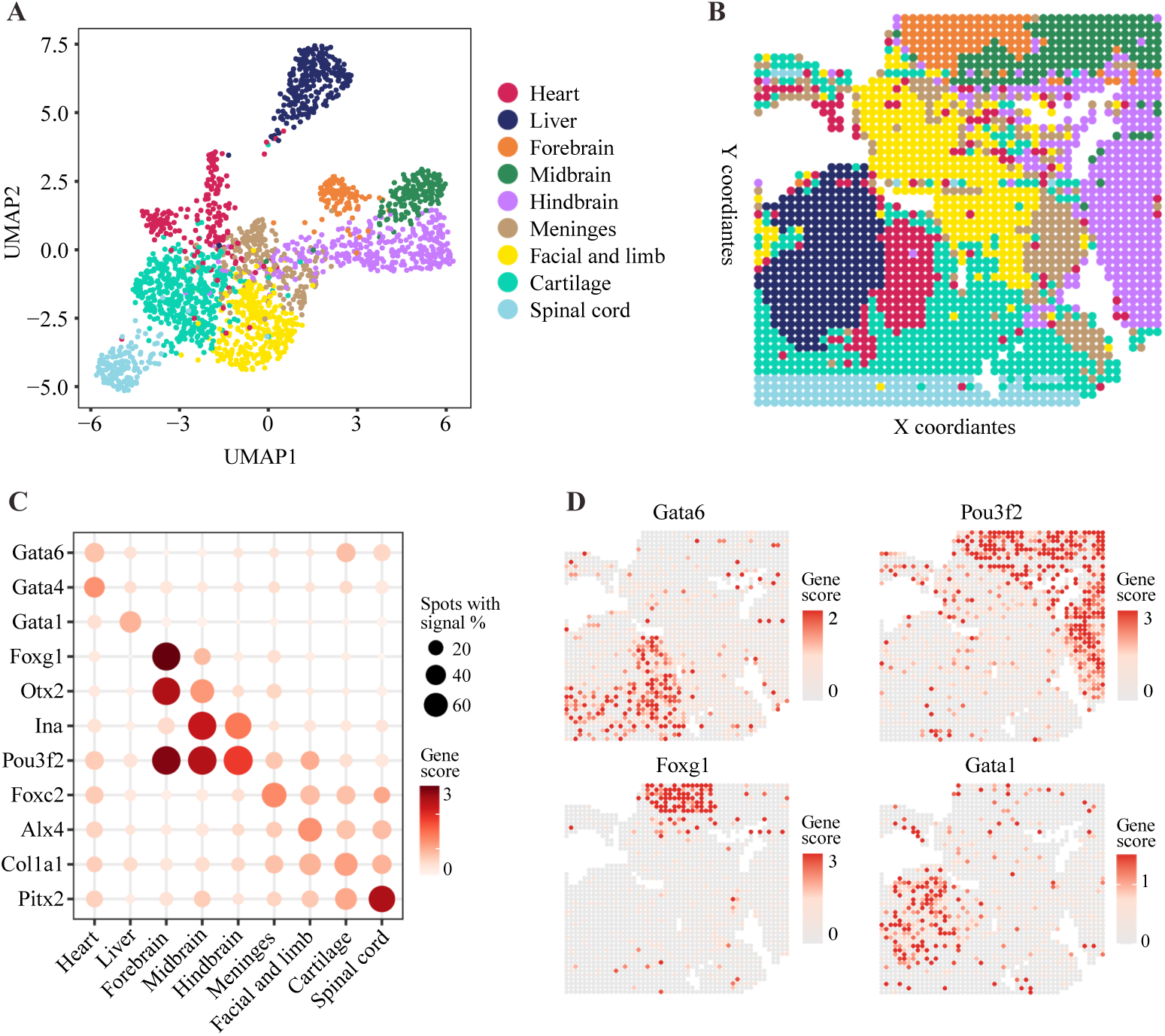
Clusters annotation based on genome-wide H3K4me3 profiles. **Related to Figure 1**. (**A-B**) UMAP and spatial distribution of clusters identified by clustering analysis. Each cluster is represented by a different color. **(C)** Clusters specific H3K4me3 signals for representative marker genes. (**D**) Spatial patterns associated with representative genes.

**Figure S2.**
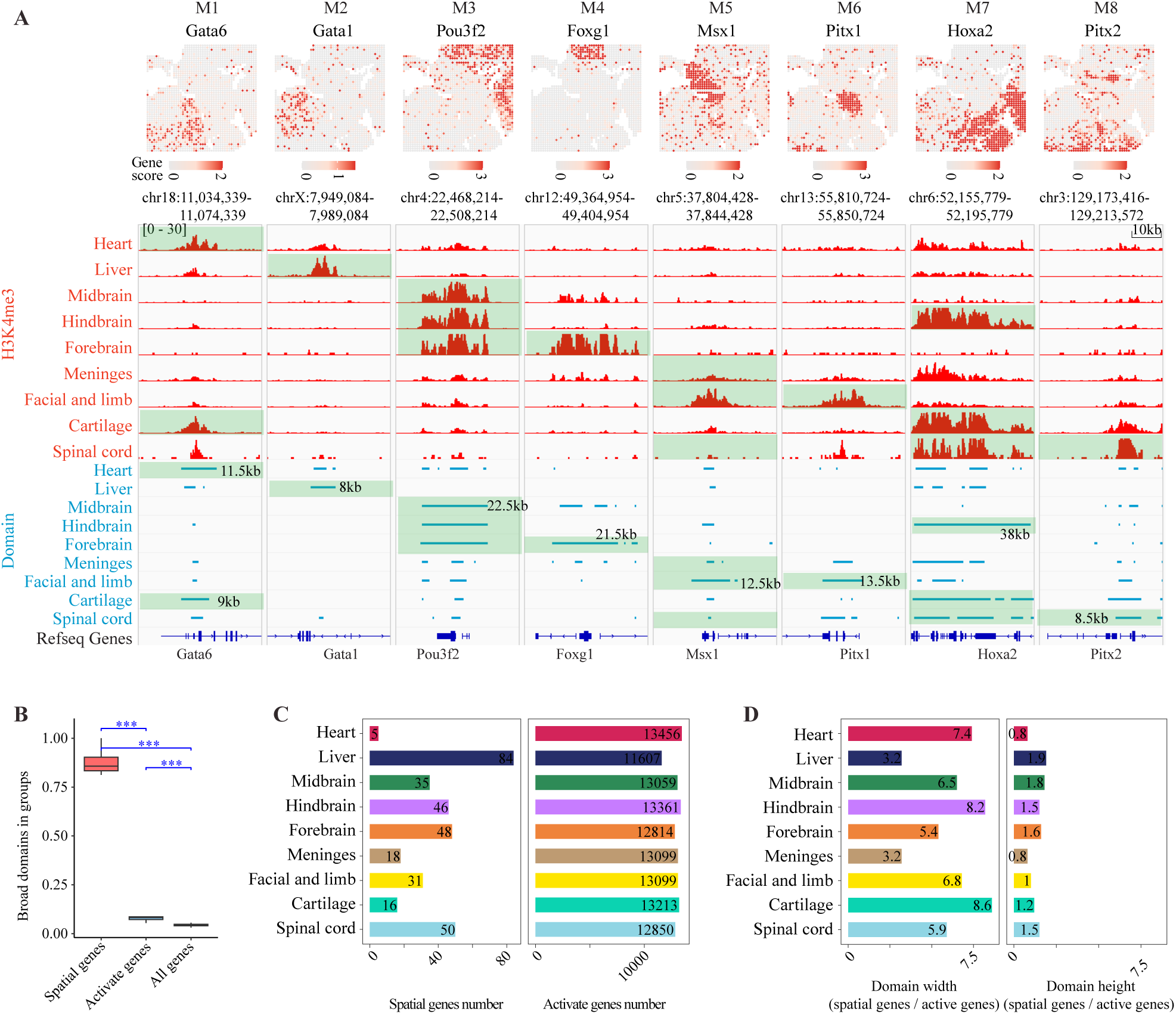
Broad H3K4me3 domains are generally associated with spatial genes. **Related to Figure 2**. (**A**) Spatial distribution, genome browser density, and domain width of spatial genes in modules. Colors on heatmap represent the H3K4me3 gene scores. Green shadows represent the modules related clusters. (**B**) Comparisons of the H3K4me3 domain enrichment between spatial genes, activate genes, and all genes (genome-wide). Y axis is the number of broad domains divided by the number of genes in each cluster. (**C**) Barplots showing the number of spatial genes and activate genes within each cluster. (**D**) Barplots showing the ratio of domain width and height between spatial genes and activate genes.

**Figure S3.**
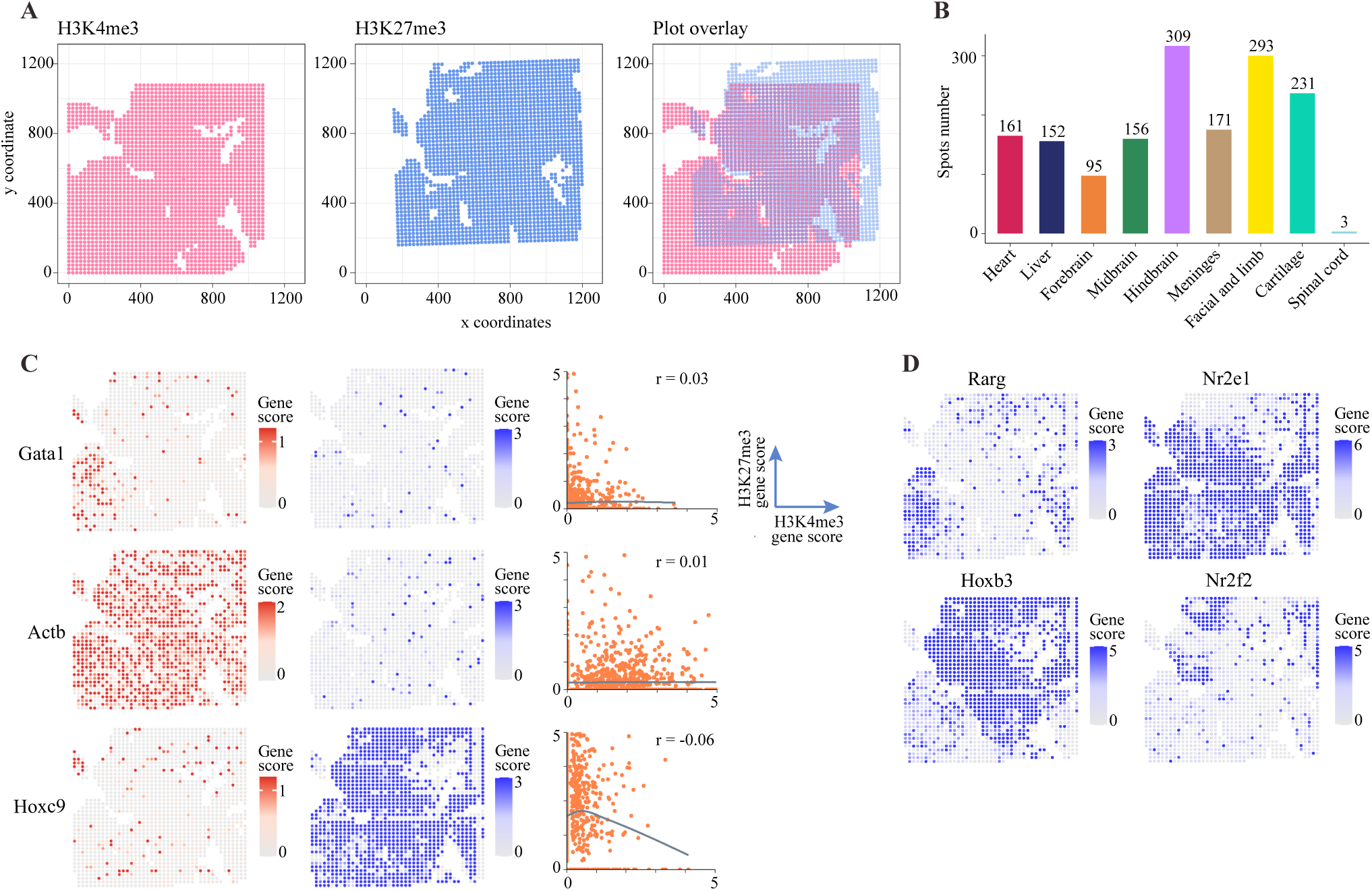
Alignment between H3K4me3 and H3K27me3, and selected genes of H3K27me3. **Related to Figure 3**. (**A**) Overlay of the aligned H3K27me3 to H3K4me3 signals. (**B**) The number of spots kept after alignment for each cluster as in Figure 1B. (**C**) Dot plot showing the H3K27me3 signals for representative marker genes as in Figure 1C. (**D**) Spatial distribution heatmaps of H3K27me3 signals of representative spatial genes.

**Figure S4.**
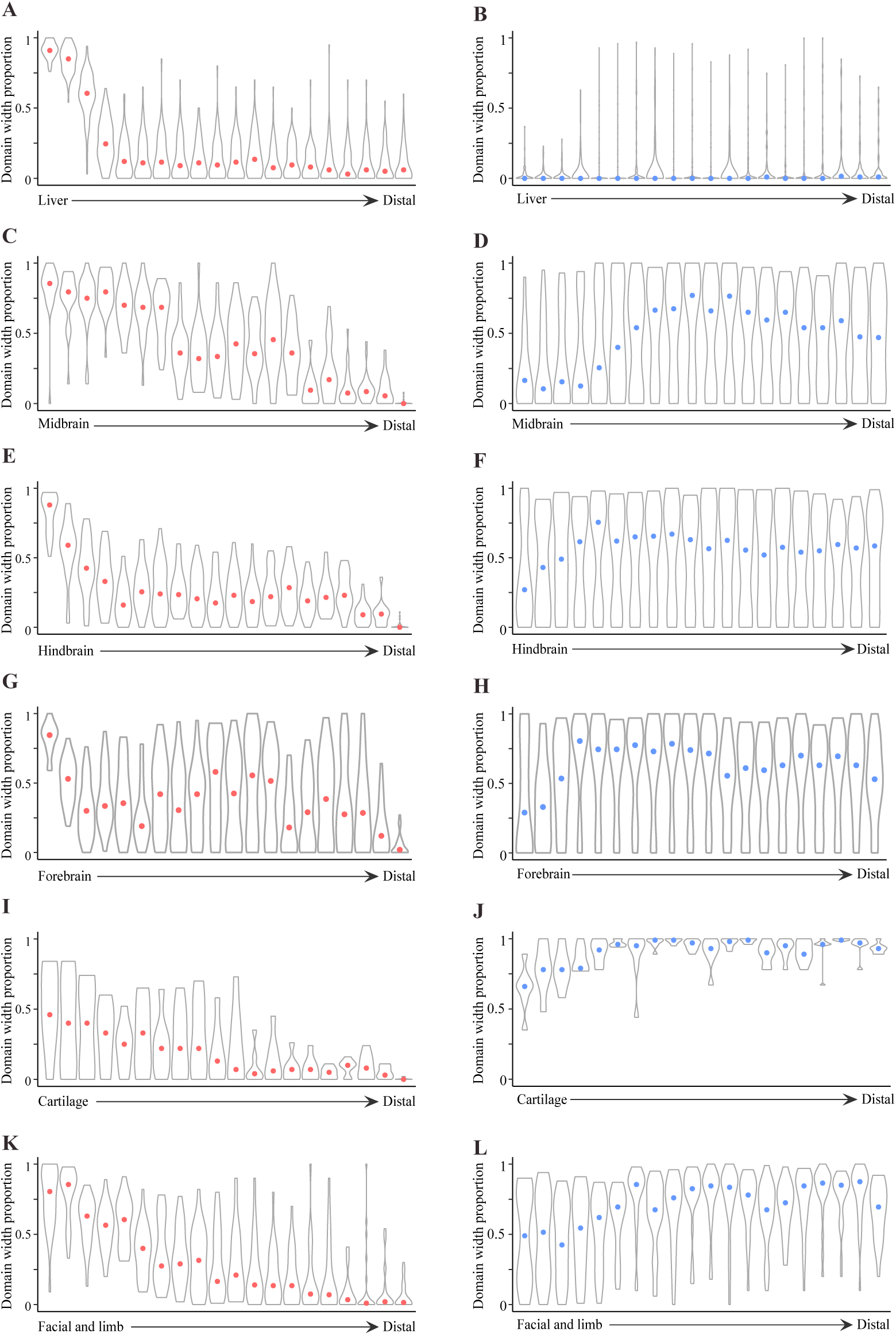
Spatial transition of chromatin states across tissues. **Related to Figure 4**. (**A-B**) The distribution of H3K4me3 and H3K27me3 (B) domain width for all spatial genes in the M2 H3K4me3 module across liver. (**C-D**) Similar to (A-B) but for M3 H3K4me3 module across midbrain. (**E-F**) Similar to (A-B) but for M3 and M7 H3K4me3 modules across hindbrain. (**G-H**) Similar to (A-B) but for M3 and M4 H3K4me3 modules across forebrain. (**I-J**) Similar to (A-B) but for M1 and M7 H3K4me3 modules across cartilage. (**K-L**) Similar to (A-B) but for M5 and M6 H3K4me3 modules across facial and limb.

